# It takes two (Ku) to repair: Mechanistic insights into the minimal NHEJ system of *Mycobacterium Tuberculosis*

**DOI:** 10.64898/2025.12.16.694593

**Authors:** Florian Morati, Evgeniya Pavlova, Anusha Budida, Selma Fornander, Elin Persson, Jagadeesh Sundaramoorthy, Raphael Guerois, Fredrik Westerlund

**Affiliations:** Department of Life Sciences, Chalmers University of Technology, Gothenburg, SE, 41296, Sweden; Institute for Integrative Biology of the Cell (I2BC), Commissariat à l’Energie Atomique, CNRS, Université Paris-Sud, Université Paris-Saclay, Gif-sur-Yvette, France

## Abstract

In this study, we used a combination of structural simulations, ensemble assays and single molecule analysis to shed light on Non-Homologous End-Joining (NHEJ) in *Mycobacterium Tuberculosis* (*Mtb*), executed by the homodimeric Ku and Ligase D (LigD). We used a monomeric form of the Ku protein to confirm the necessity of homodimerization of Ku to bind DNA. We demonstrated that *Mtb* Ku and shows limited translocation on DNA and primarily instead stays bound at the DNA ends. The reconstitution of an active *Mtb* NHEJ machinery, consisting of Ku and LigD, allowed us to characterize the dynamics of each step of the assembly of the *Mtb* NHEJ machinery at the single molecule level, revealing competition between Ku and LigD for DNA binding. *In silico* investigations highlighted key residues in *Mtb* LigD – Ku complex formation. We conducted a mutational analysis of these residues in the polymerase and the ligase domains, respectively. This investigation demonstrated that, although formation of the NHEJ complex is preserved in LigD mutants, mutations in the polymerase domain result in reduced enzymatic activity, whereas mutations in the ligase domain led to its enhancement. Finally, we present a model for NHEJ in *M. tuberculosis,* where Ku must stay tightly at the DNA ends to properly recruit and regulate LigD enzymatic activity.

**GRAPHICAL ABSTRACT:** 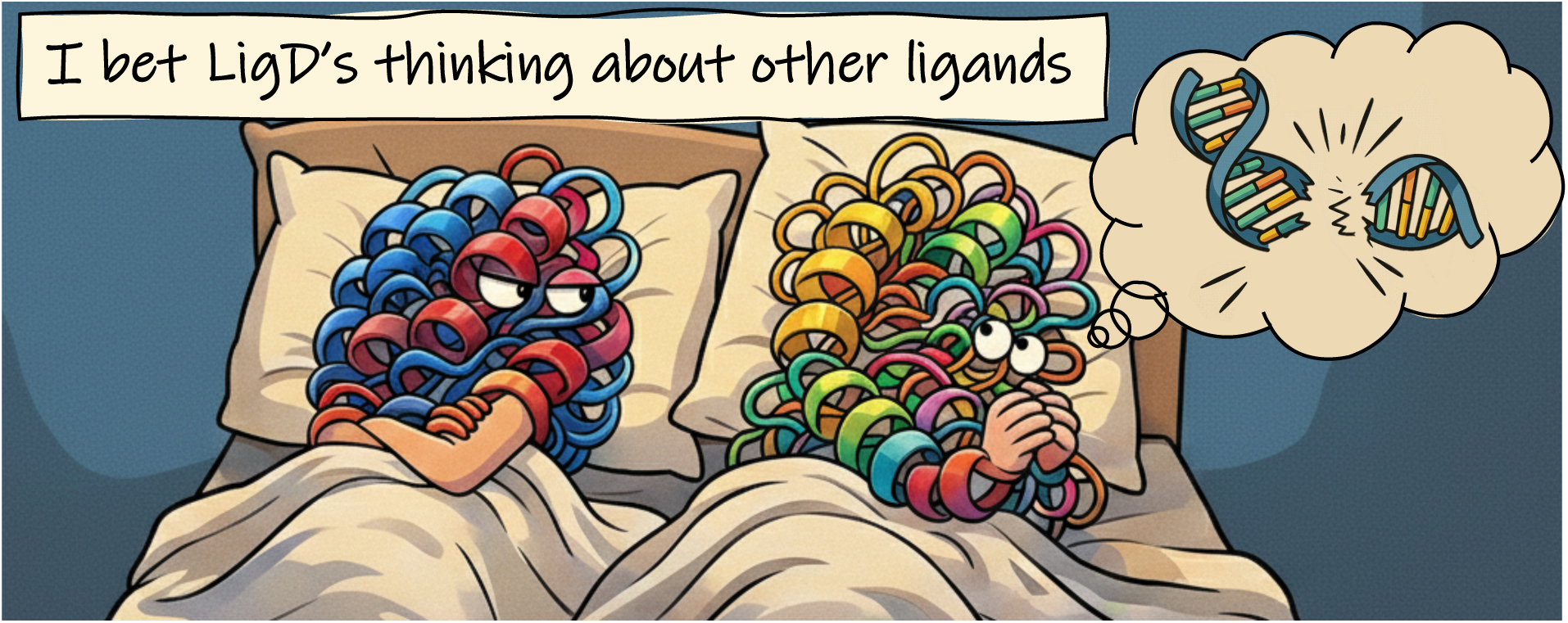

*Mtb* Ku and LigD compete for DNA binding, while interactions between Ku and the LigD polymerase (POL) and ligase (LIG) domains differentially modulate enzymatic activity. This image was generated with the assistance of ChatGPT and Gemini and subsequently edited by the authors.

## INTRODUCTION

Tuberculosis (TB), caused by *Mycobacterium tuberculosis* (*Mtb*), remains a major global health challenge. Despite being preventable and curable, more than 10 million people develop TB each year. TB has likely reemerged as the world’s leading infectious cause of death, driven in part by the spread of multidrug-resistant (MDR), pre-extensively drug-resistant (Pre-XDR), and extensively drug-resistant (XDR) strains. Understanding the genetic mechanisms that accelerate resistance evolution is therefore of critical importance^1,2^.

In other bacterial pathogens, mutator strains arising from defects in DNA repair accelerate adaptation under antibiotic pressure^3^. Although *Mtb* genomes are generally stable, the bacterium faces constant DNA damage *in vivo*, primarily from reactive oxygen and nitrogen intermediates produced by host immune cells^4,5^. DNA double-strand breaks (DSBs) are especially lethal, and during non-replicating phases of infection, when homologous recombination is unavailable, DSB repair depends exclusively on non-homologous end joining (NHEJ). Mycobacteria encode a streamlined NHEJ system composed of Ku, a homodimeric DNA-end-binding protein, and LigD, a monomeric multifunctional ligase with polymerase and phosphoesterase/nuclease activities. The presence of NHEJ protein factors in *Mtb* raises questions about the role of DSB repair, and especially Ku and LigD, in TB. Ku and LigD form a ternary complex with DNA ends, where Ku recruits and stabilizes LigD to mediate end processing and ligation. Genetic studies demonstrate that loss of either protein severely compromises NHEJ efficiency^6–8^. Beyond its role in DNA repair, NHEJ may influence *Mtb* genome evolution. Repair infidelity under genotoxic stress could increase mutagenesis, thereby accelerating the acquisition of drug resistance. This raises the possibility that defects or variations in DSB repair contribute to the emergence of highly resistant *Mtb* strains in clinical settings.

Here, we investigate NHEJ in *Mtb* , focusing on the mechanistic requirements of Ku and LigD to repair DSBs. We show that *Mtb* Ku must homodimerize in solution prior to binding to DNA ends. Upon loading on dsDNA, *Mtb* Ku stays at the DNA ends and one homodimer per DNA end is the minimal requirement for *Mtb* LigD recruitment and end-joining activity. The predicted structural models of *Mtb* Ku and *Mtb* LigD were used to design *Mtb* LigD point mutants relevant in the *Mtb* NHEJ protein complex formation. The different *Mtb* LigD mutants highlight the pivotal role of specific residues in distinctive domains of the protein, showing a prominent role of the *Mtb* LigD polymerase domain in NHEJ activity with no significant effect on the *Mtb* NHEJ complex formation in absence of DNA, pointing towards a Ku-dependent regulation of the enzymatic activity of LigD.

## MATERIAL AND METHODS

### Plasmid cloning

The His-*Mtb* Ku plasmid was created by the Protein Expertise Platform of Umeå University, by subcloning *Mtb* Ku into pET-His1a plasmid using 5’NcoI and 3’XhoI restriction sites.

The His-SNAP-*Mtb* Ku plasmid was created by the Protein Expertise Platform of Umeå University, by subcloning *Mtb* Ku into pET24d plasmid using 5’NcoI and 3’HindIII restriction sites.

The RFP-*Mtb* Ku plasmid was created by subcloning *Mtb* Ku into pRFP (Genscript) using 5’XhoI and 3’BamHI restriction sites.

The MBP-*Mtb* LigD plasmid was created by subcloning *Mtb* LigD into MBP-parallel1 (Addgene, Plasmid #134288) using 5’BamHI and 3’HindIII restriction sites.

The RFP-*Mtb* LigD plasmid was created as for RFP-*Mtb* Ku.

### Recombinant protein expression

The His-SNAP-*Mtb* Ku construct was transformed in Rosetta/pLysS bacterial cells (Novagen Cat # 70956) and induced to express the recombinant protein overnight at 15°C after addition of IPTG and pelleted. Pelleted cells were lysed in one volume of 2X lysis buffer (600 mM NaCl, 40 mM Tris pH 8, 34µg/mL benzamidine hydrochloride), supplemented with 1 mM PMSF, 2 mg Lysozyme (ThermoScientific, #89833) and a protease inhibitor cocktail (Pierce Cat #A32965). The lysate was sonicated and clarified by centrifugation at 20,000 x g for 30 min and the His-SNAP-*Mtb* Ku was recovered by chromatography over 5 ml HisTrap Excel column (Cytiva, # 17371205) pre-equilibrated in Buffer NiA (300 mM NaCl, 20 mM Tris pH 8, 1 mM EDTA) and eluted by raising the imidazole concentration to 0.5 M. The eluted fraction was concentrated using 30 kDa MWCO Ultra Centrifugal Filter (Amicon, #UFC5030) in Buffer S (300 mM NaCl, 20 mM Tris pH 8), and loaded on a HiLoad 16/600 Superdex 200 pg SEC column (Cytiva, #28989335). Estimation of molecular weight (MW) of the proteins was done using a Gel Filtration Calibration Kit (Cytiva, # 28-4038-42, Ferritin 440kDa/55mL, Canalbumin 75kDa/70mL, Ovalbumin 44kDa/80mL). Fractions of His-SNAP-*Mtb* Ku were supplemented with 2mM DTT and 10% Glycerol and stored in aliquots at -80°C.

The *Mtb* Ku-HIS construct was transformed in Rosetta/pLysS bacterial cells (Novagen Cat # 70956) and induced to express the recombinant protein overnight at 15°C after addition of IPTG and pelleted. Pelleted cells were lysed in one volume of 2X lysis buffer (600 mM NaCl, 40 mM Tris pH 8, 34µg/mL benzamidine hydrochloride) supplemented with 1 mM PMSF, 2 milligrams of Lysozyme (ThermoScientific, #89833) and protease inhibitor cocktail (Pierce Cat #A32965). The lysate was sonicated and clarified by centrifugation at 20,000 x g for 30 min and the *Mtb* Ku-HIS was recovered by chromatography over 5 ml HisTrap Excel column (Cytiva, # 17371205) pre-equilibrated in Buffer NiA (300 mM NaCl, 20 mM Tris pH 8, 1 mM EDTA), followed by a washing step in Buffer W (1M NaCl, 20 mM Tris pH 8, 1mM EDTA) and eluted by raising the imidazole concentration to 0.5 M. The eluted fraction was concentrated using 10 kDa MWCO Ultra Centrifugal Filter (Amicon, #UFC9010) in Buffer S (300 mM NaCl, 20 mM Tris pH 8), cleaved overnight using AcTEV™ Protease (Invitrogen, #12575023) and loaded on a HiLoad 16/600 Superdex 200 pg SEC column (Cytiva, #28989335). Fractions of *Mtb* Ku were supplemented with 2 mM DTT and 10% Glycerol, and stored in aliquots at -80°C.

The RFP-Ku protein preparation was purified as described for *Mtb* Ku-HIS, except no cleavage by TEV protease was done.

The MBP- *Mtb* LigD-HIS construct was transformed in Rosetta/pLysS bacterial cells (Novagen Cat # 70956) and induced to express the recombinant protein overnight at 15°C after addition of IPTG and pelleted. Pelleted cells were lysed in one volume of 2X lysis buffer, supplemented with 1 mM PMSF, 2 mg Lysozyme (ThermoScientific, #89833) and a protease inhibitor cocktail (Pierce Cat #A32965). The lysate was sonicated and clarified by centrifugation at 20,000 x g for 30 min and the MBP- *Mtb* LigD-HIS was recovered by chromatography over 5 ml MBPTrap™ HP (Cytiva, #28918779) pre-equilibrated in Buffer NiA and eluted by raising the maltose concentration to 20 mM. The eluted fraction was concentrated using 10 kDa MWCO Ultra Centrifugal Filter (Amicon, #UFC9010) in Buffer S (300 mM NaCl, 20 mM Tris pH 8), and cleaved overnight using AcTEV™ Protease (Invitrogen, #12575023) and loaded on a HiLoad 16/600 Superdex 200 pg SEC column (Cytiva, #28989335). Fractions of *Mtb* LigD were supplemented with 2 mM DTT and 10% Glycerol, and stored in aliquots at -80°C.

The RFP-LigD protein preparation was purified as described for RFP-Ku.

The list of oligos sequences (5’ to 3’), used in this work can be found in the *Supplementary information*.

The SDS-PAGE gels of the recombinant proteins used in this work can be found in *Supplementary figure 4*.

### Structural predictions: AF2 and AF3

Sequences of *Mycobacterium tuberculosis* Ku and LigD were retrieved from the UniProt database under the accession IDs P9WKD9 and P9WNV3, respectively. For the AlphaFold3 (AF3)^9^ models, sequences corresponding to the Ku dimer (either alone or fused to SNAP or RFP tags) were submitted to the AF3 server along with a 20 base pair double-stranded DNA fragment. For the AlphaFold2 (AF2)^10^ models of the Ku-LigD complexes, the protein sequences were used as input for the mmseqs2 homology search program^11^ to generate multiple sequence alignments (MSA) against the UniRef30 clustered database^12^. Homologs sharing less than 25% sequence identity with their respective query or covering less than 50% of the aligned region were excluded. When multiple homologs originated from the same species, only the sequence with the highest identity to the query was retained. Full-length ortholog sequences were then retrieved and re-aligned using MAFFT^13^. AF2 models of the Ku-LigD complex were generated by fragmenting the individual domains of Ku and LigD following the strategy described in^14^ to enhance detection sensitivity. The corresponding MSAs were concatenated such that homologs from the same species were aligned in a paired manner while sequences from different species remained unpaired. Concatenated MSAs containing 622 paired and 2000 unpaired sequences were used to generate 25 structural models (five different random seeds) for each domain delimitation using a local installation of ColabFold v1.5^15^ running three iterations of the AF2 v2.3 multimer model^10^. Model quality was assessed using the ipTM and actifpTM scores reported by AF2. Molecular graphics and analyses were performed using UCSF ChimeraX^16^.

### EMSA

Fluorescently labelled DNA substrates were prepared by the following procedure: mix oligos UP and DOWN in 1:1 ratio in TE buffer (10mM Tris pH=8.0, 1mM EDTA), heat for 5 min at 95 °C and cool down to room temperature for at least 1 h. Fluorescently labelled DNA substrate and *Mtb* Ku were mixed at the indicated final concentrations in buffer L containing 50 mM Tris pH 7.5, 30 mM KCl, 30 mM NaCl, 5 mM MgCl_2_, 2 mM DTT and incubated at room temperature for 30 min. 6% glycerol was added to the reactions prior to analysis on 6% non-denaturing polyacrylamide gel in 0.5X TBE, run at 120V for 40 minutes. The gel was imaged with ChemiDoc MP (Biorad), 60 seconds exposure (Excitation source: Blue Epi, emission filter: 532/28).

The list of oligo sequences (5’ to 3’) used in this work can be found in *Supplementary information*.

### Glass deposition

Glass coverslips (22×22 mm, #1.5) and slides (25×75 mm) were cleaned as described^17,18^. Coverslips were sonicated in acetone for 30 min; slides were soaked in 2% Hellmanex III for 1 h and rinsed with MilliQ water. Coverslips were silanized in a 1:1:100 mix of ATMS:APTES:acetone for ≥2 h at room temperature. Before use, slides and coverslips were rinsed with 70% acetone and dried with nitrogen. A 4 µl sample was applied between a coverslip-slide “sandwich” and imaged immediately. For the reaction, λ-DNA (150 nM) was incubated, if indicated,with 600 nM Ku in Buffer L at 37°C for 30 min. Then, 300 nM LigD WT or buffer without proteins was added, followed by another 90 min incubation. YOYO-1 was added (1:5 dye:base ratio) and incubated for 5-10 min at 37 °C. Samples were diluted 50 times in 0.5×TBE and deposited on functionalized glass for imaging.

### Ligation assay

Indicated concentrations of *Mtb* Ku were mixed with 100 ng of 1000 bp DNA (ThermoScientific, #SM1671) in Buffer L (50 mM Tris pH 7.5, 30 mM KCl, 30 mM NaCl, 5 mM MgCl2, 5 mM DTT and 1mM ATP if mentioned) at 37°C for 30 min. LigD was added at the indicated final concentration and incubated at 37°C for 90 min. The reaction was stopped by the addition of the stop buffer (5 mg/mL Proteinase K, 2% SDS, 0.1 M EDTA) and then incubated at 37°C for 30 min. The samples were loaded on an 0.8% agarose gel and ran at 80V for 90 min in 1XTAE buffer. The gels were stained with SYBR gold (Invitrogen, #S11494), and imaged in Chemidoc MP imaging systems (Bio-Rad). The ligation percentage (Lig%) was calculated by the following equation: Lig% = (100 ∗ lig) / (lig + un − lig) where lig and un-lig represent the band intensities of ligated (> 1kb) and un-ligated products (=1kb), respectively.

### End-joining assay and competition end joining assay

Same as for the ligation assay except for the DNA substrate used, that was generated as following: A 2kb DNA fragment, with a XhoI restriction site in the middle, was generated using oligos “EJ_DNA UP” and “EJ_DNA DOWN” on a λ-DNA template (see supplementary material). The PCR product is cleaned-up using a GenElute™ PCR Clean-Up Kit (Sigma-Aldrich, #NA1020) and digested with XhoI (NEB, #R0146S). Digested products are run on an 0,8% agarose gel and the digested DNA of 1kb in size is extracted using GenElute™ Gel Extraction Kit (Sigma-Aldrich, #NA1111). For competition assay, increasing concentration of *Mtb* Ku or *Mtb* LigD were mixed with DNA in T4 buffer (NEB, #B0202S) using the same conditions as for the ligation assay. 400 units of T4 DNA ligase were added to the reaction, incubated 30 min. at RT, followed by a heat inactivation at 65°C for 10 minutes. The gel electrophoresis, staining and imaging conditions are as described for the ligation assay.

### SNAP-pulldown

For each reaction, 3 μL SNAP-Capture Magnetic Beads (New England Biolabs, #S9145S) were washed twice with 100 μL of Buffer L and resuspended in 100 μL of Buffer L. 10 μM SNAP-Ku or SNAP peptide (New England Biolabs, # P9312S) were added to the beads for immobilization and incubated in a thermo-shaker overnight at 4°C, 600 RPM. The beads were quickly washed twice with 100 μL Buffer L and resuspended in 100 μL Buffer L with 10 μM LigD WT or LigD mutants, incubated for 90 min. in a thermo-shaker at room temperature. The beads were then washed twice and resuspended in 10 μL NuPAGE™ LDS Sample Buffer (Invitrogen, # NP0007) and incubated at 95°C for 15 minutes. The supernatant was loaded on a 4-12% Bis-Tris gel and ran at 200 V for 45 min. The gel was stained with the Pierce™ Silver Stain Kit according to the standard protocol, and imaged in Chemidoc MP imaging systems (Bio-Rad).

### Data analysis

Gel electrophoresis band intensities were measured using ImageJ. Statistical analysis was done using a paired t-test and visualized using the Graphmatik software.

## RESULTS

### Mtb Ku binds to dsDNA as an obligate homodimer

Ku binding to a DNA-end constitutes the critical initial step of NHEJ. We first evaluated the requirements of this process in *Mtb* to delineate the mechanism by which Ku is loaded onto DNA-ends. To fluorescently visualize this early NHEJ step, we purified a protein construct comprising an N-terminus (N-ter) RFP TEV-cleavable *Mtb* Ku fusion protein. Following size exclusion purification, RFP-Ku was eluted in a single peak corresponding to a theoretical molecular weight (MW) of 59 kDa, consistent with the monomeric form of the protein. This elution profile contrasts with that of the SNAP-Ku fusion protein, which exhibits a greater Stokes’ radius, and elutes in a manner consistent with a dimeric state (Fig. 1a). This difference in dimerization due to the different fusion tags was unexpected, and structural predictions were used to potentially give insight on this.

**Figure 1:**
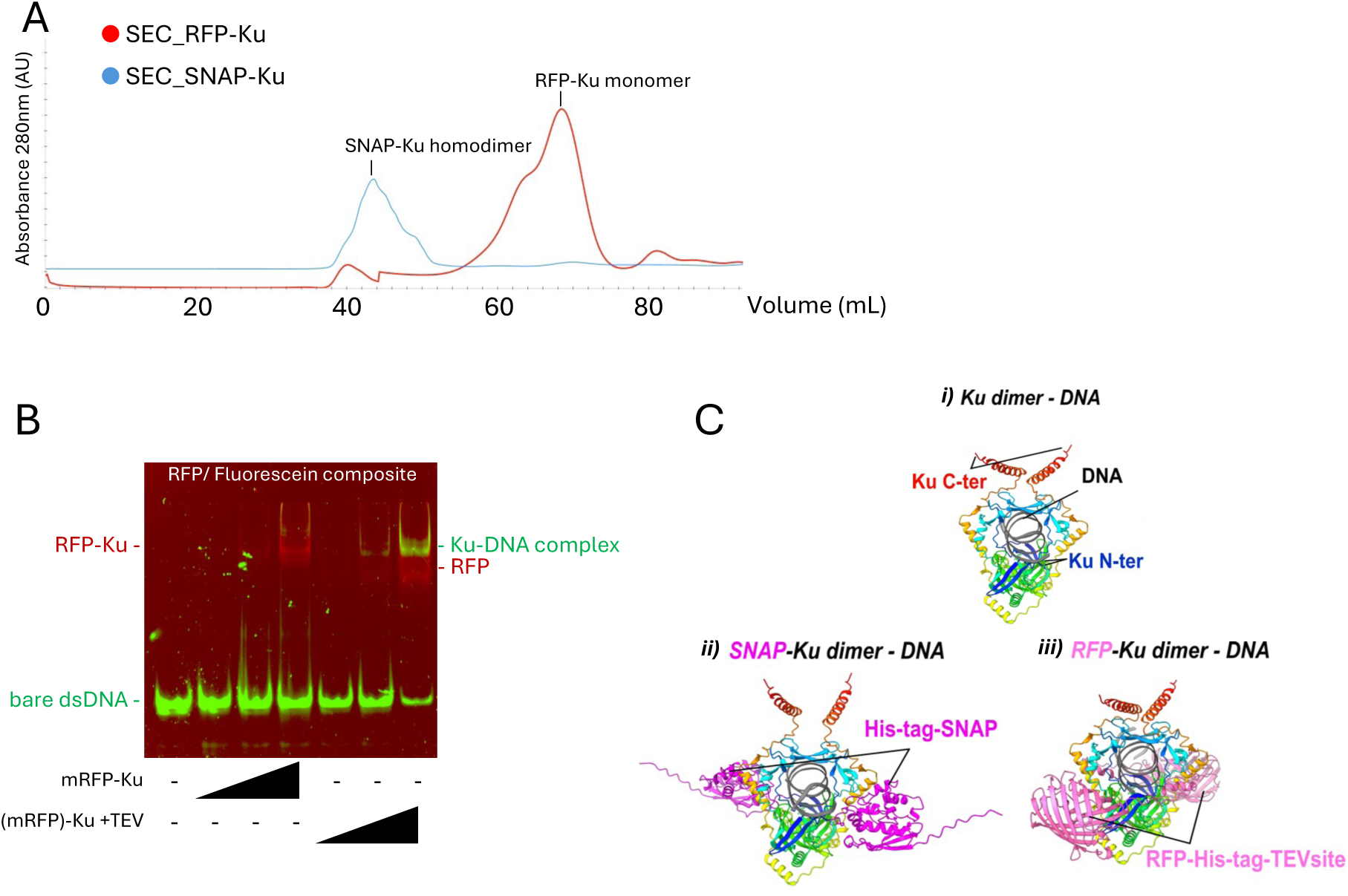
*Mtb* Ku homodimerization is required for DNA binding. a) Superposition of size exclusion profiles of SNAP-Ku (in blue) and RFP-Ku (in red) showing a difference in dimerization propensity. Estimation of proteins molecular weight (MW) was done using a Gel Filtration Calibration Kit HMW (Ferritin 440kDa/55mL, Canalbumin 75kDa/70mL, Ovalbumin 44kDa/80mL). b) *Mtb* Ku homodimerization is hindered by the N-ter RFP fusion tag. EMSA gel of DNA with increasing ratio of *Mtb* Ku protein. A FAM-labelled 30 bp/20 dT dsDNA substrate was incubated with 4, 8 and 16 *Mtb* Ku monomers/DNA molecule, prior and after TEV cleavage of the RFP tag. Reactions were analyzed on a 6% Acrylamide:Bis acrylamide gel. FAM labelled DNA was imaged in the green channel and RFP in the red channel. c) Structural models of *M. tuberculosis* Ku dimer-DNA complexes and tagged variants. AlphaFold3 (AF3) models of the *M. tuberculosis* Ku dimer bound to DNA are shown. i) Native Ku dimer-DNA complex, highlighting the N-terminal (Ku N-ter, blue) and C-terminal (Ku C-ter, red) of Ku and the bound DNA (gray). ii) Model of the His-tag-SNAP-Ku dimer-DNA complex with the His-tag-SNAP fusion protein shown in magenta. iii) Model of the RFP-His-tag-TEVsite-Ku dimer-DNA complex, with the RFP fusion shown in pink. The Ku dimer and DNA are colored as in i).

*Mtb* Ku has been described as an obligate homodimer in solution^6,19^. The RFP-Ku monomer therefore made it possible to investigate if: 1) *Mtb* Ku monomers can bind to DNA, 2) *Mtb* Ku monomers can dimerize upon DNA binding and 3) *Mtb* Ku homodimerization can occur in solution upon cleavage of the RFP-tag. These three hypotheses were investigated in an EMSA using FAM-labelled dsDNA and RFP-Ku before and after TEV treatment. Monomeric RFP-Ku did not show any DNA binding, but TEV cleavage of the RFP tag completely restored the ability of *Mtb* Ku to bind to DNA at the same level as homodimeric *Mtb* Ku alone (Fig. 1b). Hence, *Mtb* Ku needs to homodimerize to form the ring that binds to double-stranded DNA (dsDNA) ends, allowing the recruitment of other NHEJ factors^20^. Further structural work involving N-ter modified *Mtb* Ku must be done to understand the discrepancy between RFP-Ku and SNAP-Ku and elucidate which residues or motifs that are involved in homodimerization. It was first suggested that a steric hindrance form the RFP tag itself may affect the homodimerization of *Mtb* Ku. In that regard, AF3 was used to generate an RFP-*Mtb* Ku homodimer in presence of DNA (Fig. 1c). However, the presence of the TEV cleavage site at the N-ter of the *Mtb* Ku monomers is proposed to interfere with proper dimer formation, likely because this hydrophobic segment may engage the dimerization interface under our experimental conditions. This interpretation is consistent with the observation that SNAP-*Mtb* Ku, which lacks the N-ter TEV cleavage site, is able to dimerize.

### Multiple Mtb Ku can bind to dsDNA

Several *Mtb* Ku homodimers can bind to DNA ends with a homodimer footprint of 15-33 bp^6,21^. To characterize this specific DNA binding activity of *Mtb* Ku, we used an EMSA with dsDNA substrates ranging from 32 bp to 300 bp. Only one *Mtb* Ku:DNA complex was detected on a 32 bp DNA substrate with a 20dT 3’-overhang (32/20dT DNA), suggesting that *Mtb* Ku preferentially binds dsDNA and that a 32 bp dsDNA can only accommodate one *Mtb* Ku homodimer. 50 nM of the 32/20dT DNA substrate started to shift with the addition of 100 nM *Mtb* Ku (equivalent to 50 nM homodimers). The same binding activity was observed with a 50 bp blunt DNA substrate, but a second, retarded band, corresponding to a two *Mtb* Ku:DNA complex, appeared at 400 nM (Fig. 2). Interestingly, when comparing the mobility of 32/20dT overhang and 50 bp blunt DNA substrates upon increasing concentration of Ku homodimer, we observed the appearance of a third band (two Ku homodimers bound to DNA) prior to a complete shift of the unbound DNA substrate. Hence, we suggest a preference for a second Ku homodimer to bind to a Ku-bound DNA instead of bare DNA. Cooperative binding of bacterial Ku has been reported for other bacterial^22^ and mammalian Ku^23^. The 32/20dT overhang DNA substrate can only accommodate the binding of one Ku-homodimer and only two Ku-homodimers can be accommodated on the 50 bp blunt DNA, suggesting a Ku homodimer footprint of around 25 bp, in accordance with recent structural data^20^. In EMSAs using 100, 200 and 300 bp dsDNA, the same DNA binding trend was observed with a first retarded band appearing at a concentration of one *Mtb* Ku per DNA end. Increasing *Mtb* Ku concentrations lead to a distinctive two *Mtb* Ku:DNA band followed by an upper smear (Fig. 2), similar to observations for *Bacillus Subtilis* Ku^24^. Even if more than two *Mtb* Ku can load onto one DNA end, these *Mtb* Ku:DNA end complexes of higher order seem to be unstable and a stable two *Mtb* Ku:DNA end would represent a favorable platform for DNA end synapsis.

**Figure 2:**
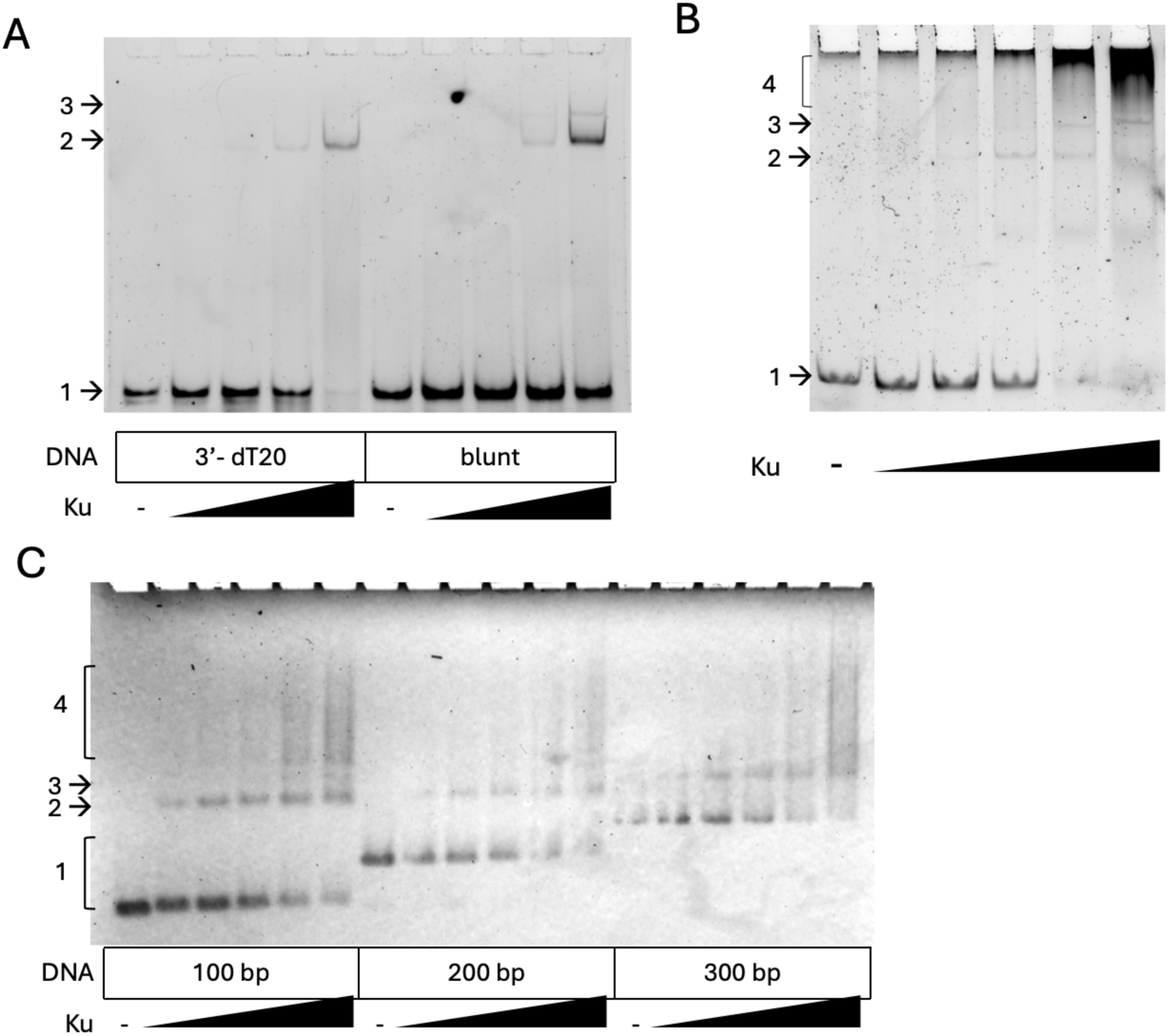
*Mtb* Ku dimers form a stable complex on dsDNA. a) *Mtb* Ku does not bind long DNA overhang/ssDNA. EMSA gel of 50 nM FAM-labelled DNA of distinctive DNA end configurations with increasing concentration of *Mtb* Ku protein. 30 bp/20 dT dsDNA and 50bp dsDNA were incubated with 50, 100, 200 and 400 nM *Mtb* Ku homodimers. The inferred compositions of DNA – *Mtb* Ku complexes from relative mobilities are noted at the side of the panel: 1) bare DNA; 2) one *Mtb* Ku homodimer:DNA; 3) two *Mtb* Ku homodimer:DNA. Reactions were analyzed on a 6% Acrylamide:Bis acrylamide gel. b) EMSA gel of 50 nM 100 bp dsDNA with increasing concentration of Ku. DNA was incubated with *Mtb* Ku at homodimeric concentrations of 0.1 to 2 μM. The inferred compositions of DNA – *Mtb* Ku complexes from relative mobilities are noted at the side of the panel: 1) bare DNA, 2) one *Mtb* Ku homodimer:DNA, 3) two *Mtb* Ku homodimer:DNA and 4) more than two *Mtb* Ku homodimer:DNA. Reactions were analyzed on a SYBR Gold post-stained 6% Acrylamide:Bis acrylamide gel. c) EMSA gel of 100-300 bp dsDNA with increasing concentration of Ku. 100 ng DNA of 100, 200 and 300 bp in size were incubated with *Mtb* Ku at homodimeric concentrations of 0.25 to 3 μM. The inferred compositions of DNA – *Mtb* Ku complexes from relative mobilities as noted at the side of the panel: 1) bare DNA, 2) one *Mtb* Ku homodimer:DNA, 3) two *Mtb* Ku homodimer:DNA and 4) more than two *Mtb* Ku homodimer:DNA. Reactions were analyzed on a SYBR Safe post-stained 2% agarose gel.

### Mtb Ku stays at the DNA ends

It has been suggested that upon binding to a DNA end, the Ku homodimer can thread along the DNA^19,21,22^. However, EMSAs with longer DNA substrates (>100 bp) highlight the low stability of more than two Ku homodimers per DNA end, making extensive Ku threading, that can be assimilated to translocation, an unlikely event for *Mtb* Ku (Fig. 2). One way to test the amplitude of *Mtb* Ku threading along DNA is to evaluate if the DNA ends are available for ligation upon *Mtb* Ku binding. We hypothesize that extensive threading of *Mtb* Ku will make the DNA ends available for enzymatic reactions. A competition end-joining assay was developed to probe this. It consists of PCR generation of a 2 kbp DNA substrate with a XhoI restriction site in the middle and non-phosphorylated DNA-ends. Upon XhoI enzyme digestion, the 1 kbp DNA substrates are purified and used for their ability to only form a linear 2 kbp DNA as a result of a complete ligation reaction. The *Mtb* Ku homodimer can bind phosphorylated as well as non-phosphorylated DNA ends as shown in EMSA experiments and ligation assays (Fig. 2 and Fig. 3a (reactions 3-5)). However, only > 11 nt phosphorylated single-stranded DNA ends can be ligated by T4 DNA Ligase (Bergstrom, NAR 2004). An increasing amount of Ku was added to 1 kbp DNA (with a one Ku homodimer:DNA end concentration increment) in T4 buffer, representing Ku-DNA incubation conditions used in standard end-joining reactions. At a concentration of one Ku homodimer per two DNA ends, *Mtb* Ku significantly interferes with T4 DNA ligase end joining activity. Addition of one Ku homodimer per DNA end leads to a complete depletion of T4 DNA ligase activity (Fig. 3a). The specific binding of *Mtb* Ku to DNA ends and its limited threading hinders the ligation of DNA ends by T4 DNA ligase. Pioneering work shows that human Ku can load and thread along dsDNA to displace histones^25^. This loading mechanistic was also demonstrated in prokaryotic NHEJ with *Bs* Ku at the single molecule level^22^ as well as in transmission electron microscopy^24^. By binding and staying at the DNA ends, *Mtb* Ku protects DNA ends from exonuclease activities, and it likely increases the probability of DNA ends meeting and promoting DNA end bridging and synapsis formation. We suggest that *Mtb* Ku homodimers can only be loaded one after another on DNA, in a train-like configuration, defined as Ku oligomerization. It was shown in electron micrograph positive staining experiments that a limited number of Ku homodimers stay at the DNA ends to promote DNA end bridging and synapsis^20^. The specific binding of Ku to DNA ends is central to its synapsis formation activity since it may increase the probability of DNA ends to meet and promote DNA ligation by LigD.

**Figure 3:**
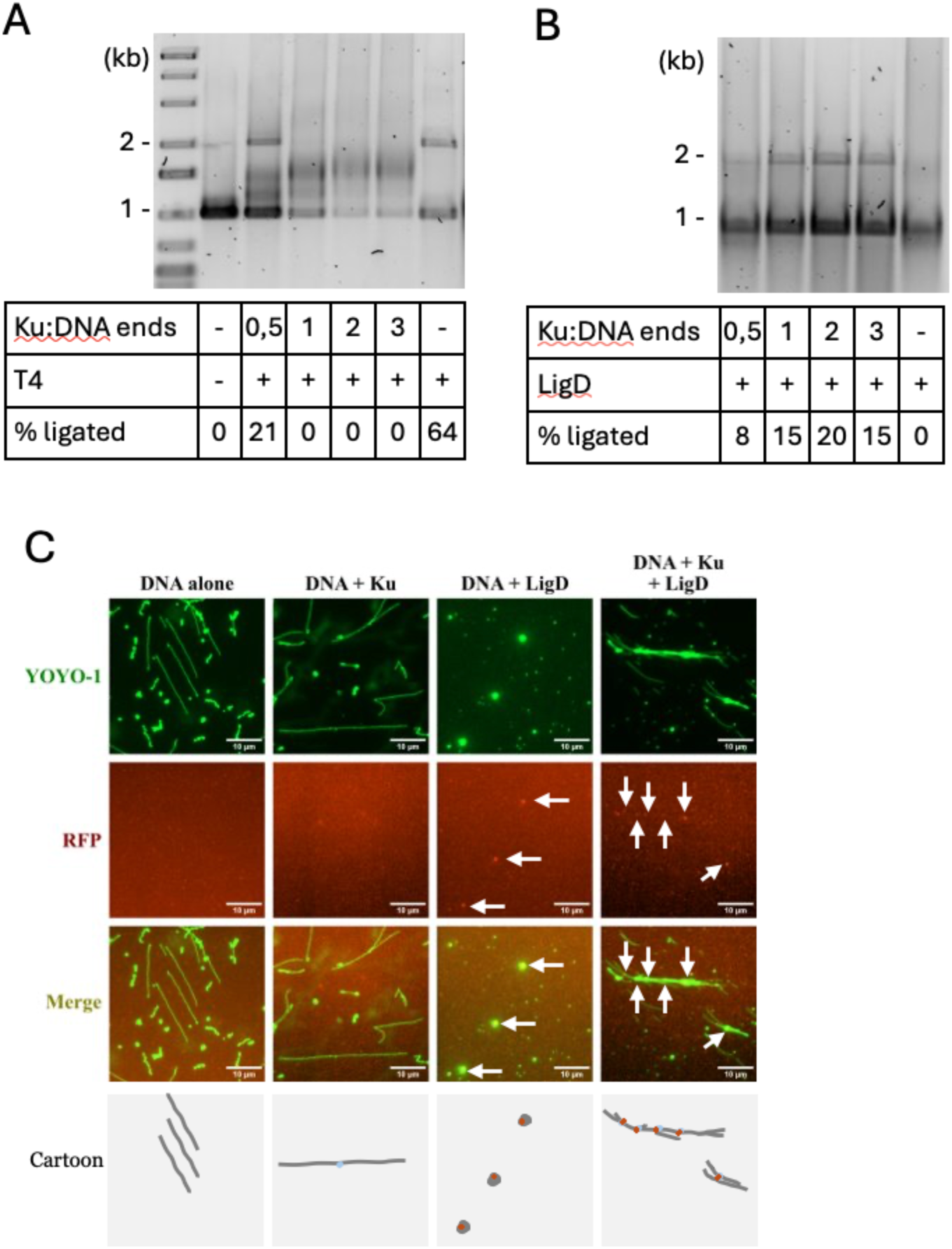
*Mtb* Ku stays at the DNA ends to orchestrate *Mtb* NHEJ. a) *Mtb* Ku impedes T4 DNA ligase activity. Increasing concentrations of homodimeric *Mtb* Ku were incubated with 1kb DNA prior to addition of 400 units of T4 DNA ligase. Reactions were deproteinated and ligation products were analyzed on a SYBR Gold post-stained 0.8% agarose gel. b) Ku-DNA complex stability is important for efficient ligation. Increasing concentration of *Mtb* Ku was incubated with 1 kb DNA prior to addition of one *Mtb* LigD/DNA-end. A ratio of two homodimeric *Mtb* Ku/DNA-end allowed the highest ligation efficiency. Reactions were deproteinated and ligation products were analyzed on a SYBR Gold post-stained 0.8% agarose gel. c) *Mtb* Ku catalyzes DNA synapsis formation allowing proper *Mtb* LigD ligation activity. DNA-protein complexes were formed using a concentration of two homodimeric *Mtb* Ku/DNA-end and/or one *Mtb* LigD/DNA-end, incubated with λ-DNA, stained with YOYO-1. The DNA-protein complexes were deposited on functionalized glass slides and imaged by fluorescence microscopy. RFP-LigD proteins are indicated by white arrows. Scale bar is 10 μm. Cartoon illustrate DNA in dark grey, *Mtb* Ku in light blue and RFP-LigD in dark orange.

### Mtb Ku orchestrates Non-Homologous End-Joining in Mtb

As previously described, *Mtb* Ku recognizes broken DNA ends and initiates the repair of the DNA double-strand breaks^21^. To better characterize the requirements of *Mtb* Ku loading to DNA ends for subsequent steps of DNA end-joining catalyzed by LigD, we performed an end-joining assay using purified *Mtb* Ku homodimer and *Mtb* LigD with the same 1 kb DNA substrate as in the competition end-joining assay. A concentration of one *Mtb* Ku homodimer bound to a DNA end was enough to recruit *Mtb* LigD and catalyze significant DNA end-joining activity. However, a concentration of two *Mtb* Ku bound to a DNA end lead to the highest DNA end-joining activity. Higher concentrations of *Mtb* Ku did not improve the *Mtb* LigD DNA end-joining activity. In the absence of DNA-bound *Mtb* Ku, no DNA-end joining products were generated, indicating that our *Mtb* LigD preparation was unable to catalyze DNA-end joining on its own, whereas the complete *Mtb* NHEJ complex was required for this activity (Fig. 3b). We determined that *Mtb* DNA ligation plateaus at concentrations of two Ku homodimers and one *Mtb* LigD per DNA end. This offers a stable protein-DNA complex allowing efficient DNA bridging, synapsis and proper recruitment of *Mtb* LigD, finally resulting in effective DNA-end joining. We hypothesize that these concentrations of *Mtb* NHEJ factors are optimal, but can vary slightly depending on the type of DNA break and DNA end accessibility. As previously mentioned, the role of *Mtb* Ku oligomerization may play a prominent role in end-joining activity. It would be interesting to produce a *Mtb* Ku mutant lacking oligomerization activity on DNA, that will allow to investigate if one *Mtb* Ku homodimer bound to DNA end is enough to keep a sufficient NHEJ activity *in vivo*.

### Mtb NHEJ activity depends on LigD adenylation

*Mtb* LigD has been demonstrated to be an ATP-dependent ligase^6^. However, in end-joining assays employing a ratio of two *Mtb* Ku proteins and one *Mtb* LigD protein per DNA-end, performed in ligation buffer either containing or lacking ATP, we observed comparable ligation efficiencies irrespective of ATP presence in the reaction mixture (Fig. S1). We cannot exclude the possibility that our *Mtb* LigD preparations become self-adenylated within the host cells during expression, or that residual ATP present in the preparations is sufficient to support DNA end joining, thereby accounting for this observation. Self-adenylation of *Mtb* LigD remains the most likely explanation in our case. In principle, unadenylated *Mtb* LigD would release pyrophosphate during the adenylation step, which could serve as an indirect readout of ligation activity if adenylation occurs under the assay conditions^20,21^.

### Mtb Ku catalyzes the pairing of DNA ends allowing end-joining activity by Mtb LigD

Once the *Mtb* Ku homodimer binds to a DNA end, it catalyzes DNA end bridging, synapsis and recruitment of *Mtb* LigD to initiate the repair process by NHEJ. To decipher the timeline of these events, it is essential to apply single molecule techniques, allowing the visualization of discreet populations that constitute intermediate protein-DNA complexes. To do this, we used a glass deposition assay^17,18^. Here, λ-DNA, which has 12 nts 5’complementary overhangs, was mixed with *Mtb* Ku at concentration corresponding to two *Mtb* Ku homodimers per DNA end, and the DNA-protein complexes were deposited on silanized glass slides and imaged with a fluorescence microscope. By comparing with bare λ-DNA, we observed the appearance of a population of DNA molecules longer than a single λ-DNA molecule. We attribute this population to λ-DNA molecules forming synapses and bridges in presence of Ku, as also observed for *Bs* Ku^22^. When RFP-labeled *Mtb* LigD was added to *Mtb* Ku bound λ-DNA at a concentration of two *Mtb* Ku and one RFP-*Mtb* LigD per DNA-end, we observed RFP-*Mtb* LigD at the center of long DNA clusters (Fig. 3c). When *Mtb* LigD was added alone to λ-DNA, aggregates were formed. We propose that the latter activity is due to non-specific DNA-binding activity that may be slower than the specific binding to *Mtb* Ku on DNA-ends. The DNA binding probably has a negligeable effect at the early steps of *Mtb* NHEJ, but potentially a more important role later by clustering DNA ends, and offering protection from endonuclease activity during DNA DSBs repair. This could be a platform for recruitment of protein factors that can remove hindering DNA-bound proteins, but further investigation is needed to confirm this suggestion.

Our single molecule data indicate that the *Mtb* NHEJ factors remain associated with repaired DNA. *Mtb* LigD proteins localize to sites where two DNA molecules are joined, consistent with observations reported for *Bs* Ku^22^. These findings complement our ligation assay results, in which DNA molecules remained joined following proteinase K treatment, demonstrating successful DNA repair under the same experimental conditions. Recent work showed that multiple DNA-bound homodimeric Ku can recruit and activate UvrD-1 unwinding activity^26^. The physiological meaning of this *Mtb* Ku-UvrD1 interaction remains unclear, but the role of UvrD-1 in NHEJ seems unlikely, suggesting that multiple DNA-bound homodimeric Ku may have a more pivotal role in other repair pathways than in NHEJ.

### Mtb LigD can interfere with Mtb Ku DNA binding

Since *Mtb* LigD shows DNA binding activity, likely due to its OB fold, we investigated if *Mtb* LigD competes with *Mtb* Ku for DNA ends, using a competition end-joining assay. *Mtb* LigD interferes significantly with T4 DNA ligase activity at concentrations slightly lower than one LigD per DNA molecule and it completely disrupts T4 DNA ligase activity at a concentration equal or higher than one LigD per DNA end (Fig. S2). We then investigated if *Mtb* LigD may interfere with Ku loading to DNA ends and subsequent DNA-end joining. For that purpose, increasing concentrations of *Mtb* LigD were first added to DNA, followed by addition of Ku, incubated following the same conditions as described for the end-joining assay. Addition of *Mtb* LigD prior to *Mtb* Ku disrupts the *Mtb* end-joining activity only at LigD concentrations higher than one LigD per DNA end (Fig. S3). We propose that the formation of DNA condensates limits the availably of DNA ends to be ligated. Investigations by the Andres laboratory showed that *Mtb* Ku and LigD compete for DNA substrates, with Ku displaying a higher affinity for DNA ends, thereby likely interfering with the polymerisation activity of LigD on short DNA substrates^27^. The addition of *Mtb* LigD prior to *Mtb* Ku addition is showing a smaller effect than with T4 DNA ligase on DNA end-joining activity, which may be due to *Mtb* Ku:*Mtb* LigD protein-protein interactions that mitigate DNA condensates formation triggered by not complexed *Mtb* LigD (Fig. S3). Altogether, these observations reveal an intricate Ku–LigD interplay within the *Mtb* NHEJ complex that governs its enzymatic activity.

### Mtb LigD ligation activity is modulated by its interaction with Mtb Ku

*Mtb* LigD consists of three distinctive domains: polymerase (POL), nuclease/phosphoesterase (NUC/PE) and ligase (LIG)^28^ domain. *In silico* investigations using AF2 highlighted two potential interfaces in the POL and LIG domains of *Mtb* LigD that potentially participate in the protein-protein interaction with *Mtb* Ku in absence of DNA (Fig. 4a). Within these two interfaces of *Mtb* LigD, six individual amino acid residues were considered as critical for the interaction between *Mtb* LigD and *Mtb* Ku. Point mutations were designed to test this hypothesis, in the POL domain: D162R, V194D, R198E, and in the LIG domain: D522R, K579E and L580E (Fig. 4a). To validate the predicted alterations in the protein-protein interaction of the *Mtb* Ku and *Mtb* LigD mutants, we used an anti-SNAP tag pulldown assay using SNAP tagged *Mtb* Ku (homodimer) as bait and *Mtb* LigD WT and mutants as prey. *Mtb* LigD WT and all mutants were recovered equally well at equilibrium using this assay, showing no significant disruption in the *Mtb* Ku-*Mtb* LigD interaction caused by the point mutations in *Mtb* LigD (Fig. 4b). Hence, we suggest that complete disruption of the *Mtb* Ku-*Mtb* LigD interaction should be a combination of point mutations in the *Mtb* LigD and *Mtb* Ku interfaces. Since all *Mtb* LigD versions exhibited a similar binding for *Mtb* Ku at equilibrium, we then investigated if these *Mtb* LigD point mutations affected the ligation of dsDNA substrates. During a ligation event, dsDNA substrates were first bound at a constant concentration of two *Mtb* Ku homodimers per DNA end, followed by the addition of *Mtb* LigD WT or mutants at a concentration of one *Mtb* LigD per DNA end, in the same experimental conditions as in the previous end-joining assays (Figure 3b) except 1kb NoLimits^TM^ DNA substrate with phosphorylated blunt ends was used. Two POL domain mutants (D162R and V194D) showed a significantly reduced ligation activity compared to WT, while R198E did not show any significant reduction in ligation activity (Fig. 4c). We propose that the ligation activity of the *Mtb* LigD R198E mutant remains similar to that of the wild type because the mutation lies near *Mtb* Ku E256, which is predicted to form a salt bridge with *Mtb* LigD R198. The presence of the adjacent *Mtb* LigD R202 residue may compensate for the loss of this favorable interaction, thereby preserving ligation activity despite the R198E substitution. The POL domain of *Mtb* LigD physically associates with *Mtb* Ku, thereby stimulating polymerization and likely mediating the recruitment of bacterial NHEJ ligases to Ku-bound double-strand breaks (DSBs) *in vivo*. The retention of this domain in nearly all prokaryotic Ku-associated ligases suggests that this recruitment mechanism is evolutionarily conserved^28^. It has been demonstrated that the LIG domain has a higher affinity for DNA than the POL domain^28^. The LIG domain mutants D522R and K579E demonstrated significantly increased ligation activity compared to WT Mtb LigD (Fig.4c), likely because the reduced DNA binding affinity allows for higher turnover, similar to *Bs* LigD^22,29^. In contrast, the L580E mutant did not show this increase; it exhibited activity levels similar to WT (no significant reduction), though this specific result may be obscured by nucleolytic degradation of the substrate.

**Figure 4:**
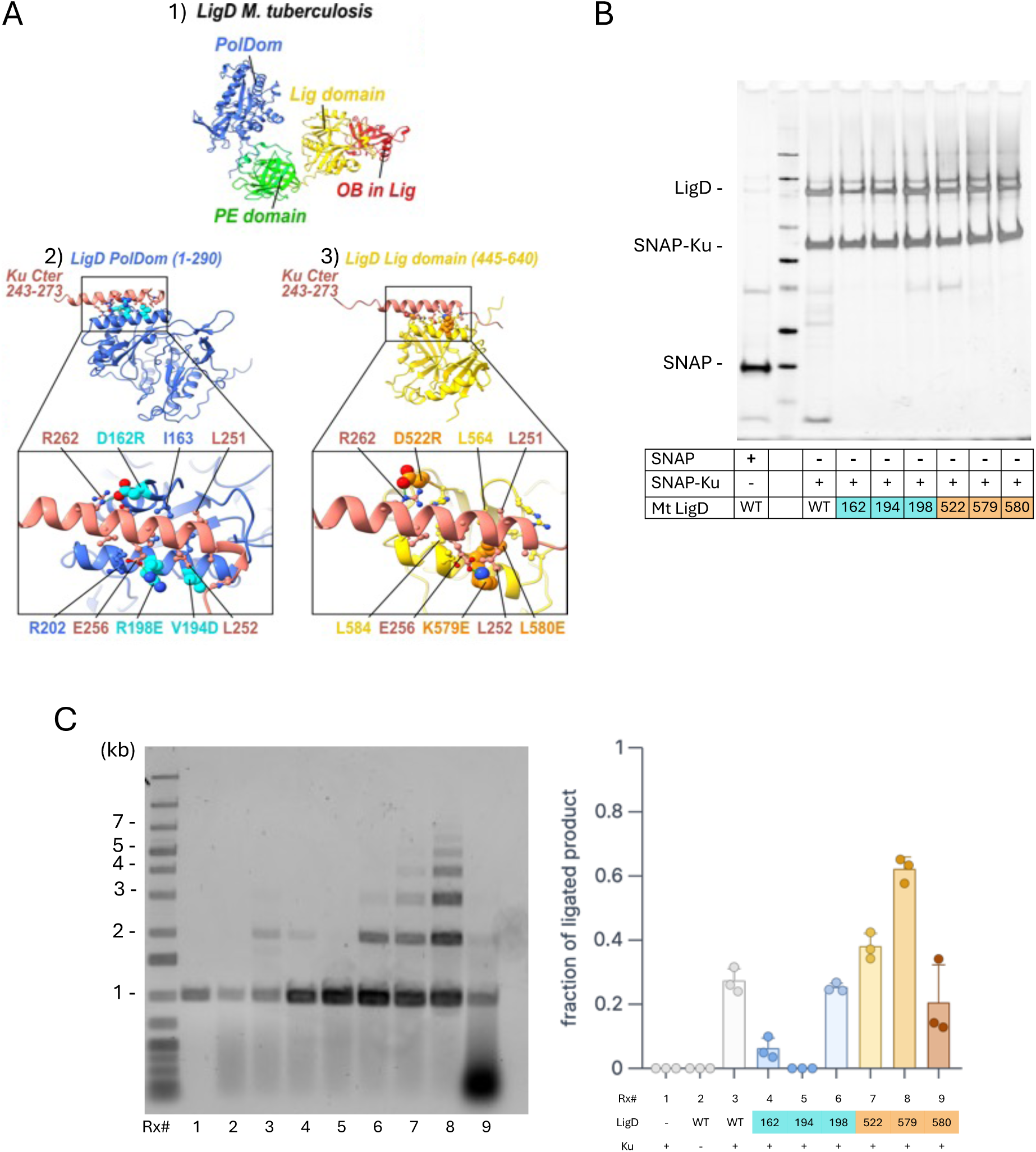
Mutants of *Mtb* LigD interact with *Mtb* Ku, and show distinctive ligation activities. a) Predicted interactions between the Ku C-terminal region and LigD domains from *M. tuberculosis*. AlphaFold2 (AF2) models of LigD and its domains. 1) Domain organization of LigD, including the polymerase domain (PolDom, blue), ligase domain (Lig, yellow) containing an OB subdomain (red), and phosphoesterase domain (PE, green). 2) Predicted interaction between the Ku C-terminal region (residues 243-273, salmon) and the LigD PolDom (residues 1-290, blue). Residues at the interface are highlighted and labeled, with interface confidence scores of AF2 ipTM = 0.87 and actfipTM = 0.86. 3) Predicted interaction between the same Ku C-terminal region and the LigD Lig domain (residues 445-640, yellow), showing key contact residues. Interface confidence scores are AF2 ipTM = 0.79 and actfipTM = 0.77. Insets in Middle and Right panels show detailed views of the Ku C-terminal helix (salmon) docking onto each LigD domain. The amino acids that were mutated are represented as large spheres and in a different color (cyan and orange for PolDom and Lig domains, respectively). b) *Mtb* LigD WT and mutants interact with *Mtb* Ku. The 6 *Mtb* LigD mutants (polymerase domain: D162R, V194D, R198E and ligase domain: D522R, K579E and L580E) were pulled down with anti-SNAP beads. Proteins were identified on a denaturing SDS-PAGE followed by a silver stain. *Mtb* LigD MW= 81kDa, SNAP-*Mtb* Ku MW= 53kDa, SNAP peptide MW= 20kDa. c) *Mtb* LigD mutants show a wide variety in ligation efficiency. *Mtb* Ku and 1kb DNA (NoLimits) were incubated at a two homodimeric Mtb Ku/DNA-end ratios followed by the addition of one *Mtb* LigD/DNA-end. Reactions were deproteinated and ligation products were analyzed on a SYBR Gold post-stained 0.8% agarose gel.

Although qualitative pull-down assays showed no detectable differences in complex formation between *Mtb* LigD mutants and *Mtb* Ku, the mutants clearly displayed variations in overall NHEJ enzymatic activity. Quantitative analysis by Microscale Thermophoresis (MST), performed by the Andres laboratory^27^, resolved this discrepancy. Consistent with findings for POL-domain mutants, the MST data demonstrated that most LIG-domain mutants actually exhibit reduced binding affinity, with the exception of K579E. Collectively, these results indicate that the strength and the topology of the Ku–LigD interaction, positively stimulates enzymatic function, leading to enhanced ligation activity.

The results from the Ku-LigD interaction studies, suggest that AF2 predictions must be complemented with functional characterization. The LigD POL domain was suggested to be pivotal (with the highest confidence level) in the *Mtb* NHEJ complex formation with DNA. This emphasizes the importance of running experimental assays to evaluate AF predictions. The *Mtb* Ku homodimer undergoes structural changes upon DNA binding which likely change the way *Mtb* LigD interacts with the *Mtb* Ku-DNA complex during ligation, and it likely affects the ability of *Mtb* LigD enzymatic domains to access DNA^19^.

To conclude, our results, in agreement with the findings of the Andres Lab^27^, indicate that *Mtb* Ku-LigD interactions play roles extending beyond the simple recruitment of *Mtb* LigD to broken DNA ends, as initially anticipated. These differences in protein-protein and protein-protein-DNA interactions underscore the tight regulation of *Mtb* NHEJ from its earliest stages. The strong and specific binding of the *Mtb* Ku homodimer to DNA ends remains essential for efficient DNA end joining and for proper recruitment and positioning of the *Mtb* LigD during DNA processing. Furthermore, interactions between *Mtb* Ku and the distinct enzymatic domains of *Mtb* LigD modulate both *Mtb* LigD activity and the overall *Mtb* NHEJ enzymatic activities (Fig.5).

**Figure 5:**
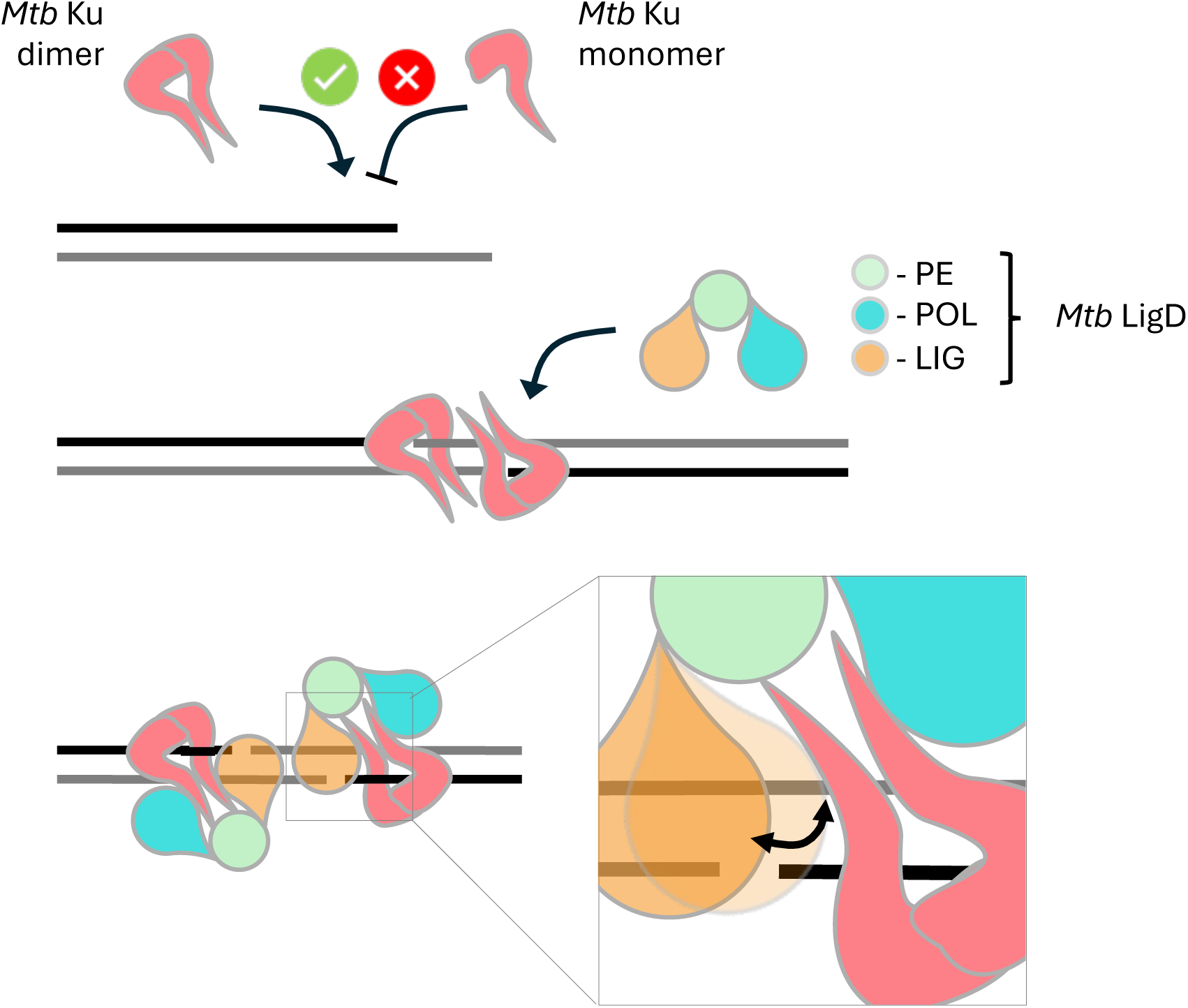
proposed model of Mtb NHEJ. Mtb Ku must dimerize to bind double-stranded DNA. These Mtb Ku dimers remain stably associated with broken DNA ends, thereby promoting DNA end joining and facilitating the recruitment of Mtb LigD. Mtb Ku interacts with both the polymerase (POL) and ligase (LIG) domains of Mtb LigD, modulating LigD’s access to DNA and consequently regulating its enzymatic activities.

## ACKNOWLEDGEMENTS

We acknowledge constructive discussions with the lab of Sara Andres at McMaster University.

The authors acknowledge the use of ChatGPT 4.0 (OpenAI) and Gemini (Google) to improve the clarity and readability of the manuscript text. Additionally, these tools were used to generate the initial concept for the graphical abstract. The authors reviewed and revised all AI-generated content and take full responsibility for the accuracy and integrity of the imagery.

## AUTHOR CONTRIBUTIONS

Florian Morati: Conceptualization, Data curation, Formal analysis, Investigation, Methodology, Visualization, Writing – original draft. Evgeniya Pavlova: Data curation, Formal analysis, Investigation. Anusha Budida, Selma Fornander, Elin Persson, Jagadeesh Sundaramoorthy: Data curation. Raphael Guerois: Data curation, Formal analysis, Resources, Software, Writing – review & editing. Fredrik Westerlund: Funding acquisition, Project administration, Resources, Supervision, Validation, Writing – review & editing.

## SUPPLEMENTARY DATA

Supplementary Data are available at NAR online.

## DATA AVAILABILITY

The data underlying this article will be shared on reasonable request to the corresponding author.

