## Supplementary figures and tables for "It takes two (Ku) to repair: Mechanistic insights into the minimal NHEJ system of *Mycobacterium Tuberculosis*"

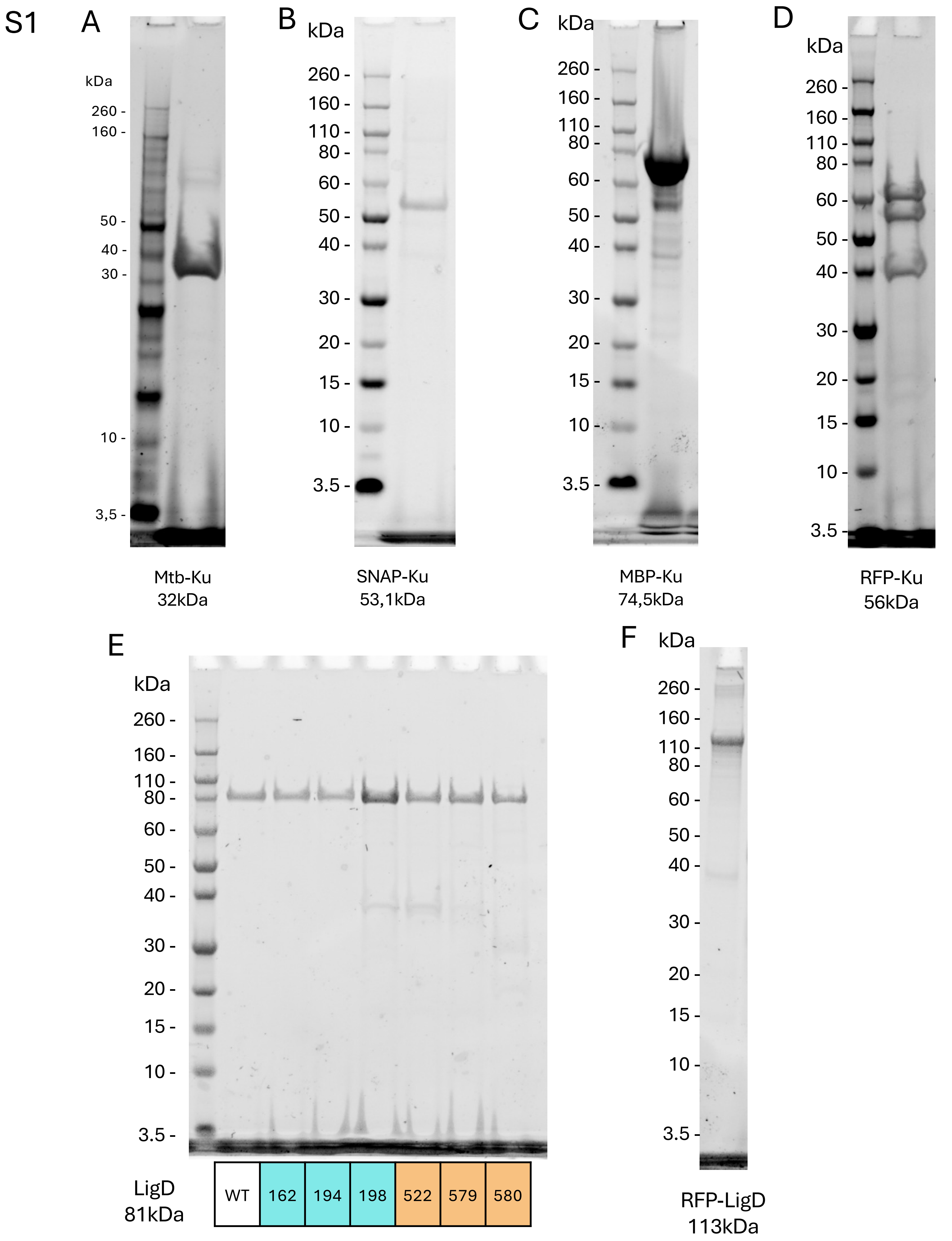


**Supplementary figure 1:** Recombinant protein preparations used in this work.

4-12% Bis-Tris gel of a) *Mtb*-Ku, b) SNAP-Ku, c) MBP-Ku, d) RFP-Ku, e) *Mtb*-LigD WT and polymerase domain: D162R, V194D, R198E and ligase domain: D522R, K579E and L580E mutants, f) RFP-LigD, stained in Coomassie blue.

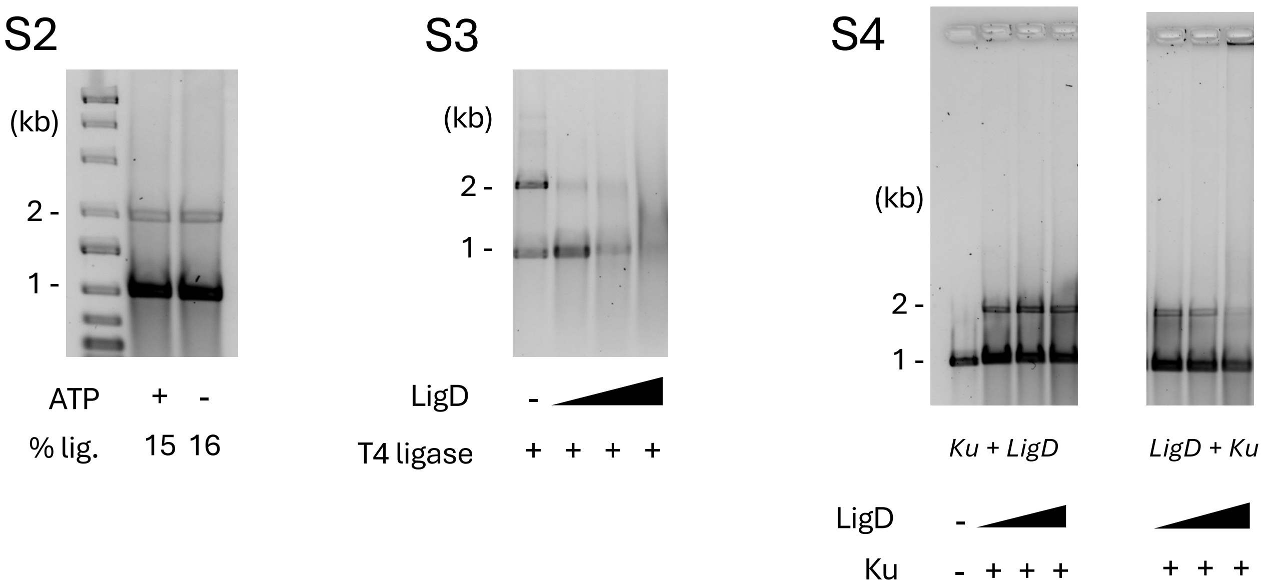


**Supplementary figures 2-4:** *Mtb* Ku and *Mtb* LigD compete for DNA substrates.

S2) Addition of ATP is not needed for efficient *Mtb* LigD ligation activity. *Mtb* Ku and 1kb DNA were incubated at a two homodimeric *Mtb* Ku/DNA-end ratio followed by the addition of one *Mtb* LigD/DNA-end, in a buffer supplemented or not with ATP. Reactions were deproteinated and ligation products were analyzed on a SYBR Gold post-stained 0.8% agarose gel.
S3) *Mtb* LigD impedes T4 DNA ligase activity. Increasing concentration of *Mtb* LigD were incubated with 1kb DNA prior to addition of 400 units of T4 DNA ligase.
S4) *Mtb* LigD impedes *Mtb* Ku binding to dsDNA. Increasing concentration of *Mtb* LigD were incubated with 1kb DNA after or prior to addition of two homodimeric *Mtb* Ku/DNA-end. Reactions were deproteinated and ligation products were analyzed on a SYBR Gold post-stained 0.8% agarose gel.

### Supplementary information

### List of oligos sequences (5' to 3'), used in this work:

Cloning *Mtb* NHEJ factors

| | **UP** | **DOWN** |
|---|---|---|
| RFP-Ku | GCCATCTCGAGATGCGCGCGATTTGGACAGGG | CATGCGGATCCTTATGGCGGAGTGGGTACATTCGAG |
| MBP-LigD | GCCATGGATCCGATGGGCTCAGCTTCGGAACAGC | GTGCCAAGCTTTCAGTGGTGGTGGTGGTGGTG |
| RFP-LigD | GCCATCTCGAGATGGGCTCAGCTTCGGAACAGC | CACAAGGATCCTCAATTCTCTTCACGAACAACTTCGGACGGTTTC |

Mutagenesis of *Mtb* LigD

| | **UP** | **DOWN** |
|---|---|---|
| D162R | CGCATTGGCCTGGTGACATTCCCG | TGCCAGCAAGTCGCGCACCGC |
| V194D | GACCTGGCCAAACGCGTAGCGC | CGTAGCGCCACGAGAACTAACCGGC |
| R198E | GAAGTAGCGCAGCGTCTGGAACAGGCG | TTTGGCCAGCACCGTAGCGCCACG |
| D522R | CGCCACCATGTCGTCCTCGATGGCGAA | GGCCAAATCTTCTGCCAGGGCGCG |
| K579E | GAACTGTTGGAAACTCTCGCTAACGCG | GCGCCGATCTTGGTAACGGGTCCCC |
| L580E | GAATTGGAAACTCTCGCTAACGCG | TTTGCGCCGATCTTGGTAACGGGTCCCC |

EMSA

| | **UP** | **DOWN** |
|---|---|---|
| 32/20dT | GCCCTGCTGCCGACCAACGATGGTTACATTCC | GGAATGTAACCATCGTTGGTCGGCAGCAGGGCTTTTTTTTTTTTTTT TTTTT |
| 50bp blunt end | GCTAAGTCATCGTCTGTCGATCGATCTCTCGAGTAGTACAGGTCT AGAGG | CCTCTAGACCTGTACTACTCGAGAGATCGATCGACAGACGATGACT TAGC |

DNA substrate for End joining assay

| | **UP** | **DOWN** |
|---|---|---|
| EJ_DNA | AGGATTCATTGTCCTGCTCAAAGTCC | CCATTTCTATACTCATCAAACTGTAGGGGTTG |
